# A shared representation of order between encoding and recognition in visual short-term memory

**DOI:** 10.1101/080317

**Authors:** Kristjan Kalm, Dennis Norris

**Affiliations:** Cognition and Brain Sciences Unit, Medical Research Council 15 Chaucer Road, Cambridge, CB2 7EF, UK

## Abstract

Most complex tasks require people to bind individual stimuli into a sequence in short term memory (STM). For this purpose information about the order of the individual stimuli in the sequence needs to be in active and accessible form in STM over a period of few seconds. Here we investigated how the temporal order information is shared between the presentation and response phases of an STM task. We trained a classification algorithm on the fMRI activity patterns from the presentation phase of the STM task to predict the order of the items during the subsequent recognition phase. While voxels in a number of brain regions represented positional information during either presentation and recognition phases, only voxels in the lateral prefrontal cortex (PFC) and the anterior temporal lobe (ATL) represented position consistently across task phases. A shared positional code in the ATL might reflect verbal recoding of visual sequences to facilitate the maintenance of order information over several seconds.

## 1 Introduction

One of the most important features of human short term memory (STM) is the ability to bind individual stimuli into a sequence. A host of complex behaviours including language processing, vocabulary acquisition, and chunk formation are thought to rely on sequence encoding in STM (see Hurlstone, Hitch, & Baddeley, 2014, for a review). For these purposes information about the order of the individual stimuli in the sequence needs to be in active and accessible form in STM over a period of few seconds (Botvinick & Watanabe, 2007; Baddeley, 2003). Research has shown that such stimulus’ position in the sequence is encoded in STM separately and independently of their identity (Henson & Burgess, 1997; Henson, Norris, Page, & Baddeley, 1996; Page & Norris, 2009, Figure 1A). From hereon we refer to such neural representation of the item’s position in the sequence as *positional code*. Figure 1C gives an example of a simple positional code showing the responses of position-sensitive neurons from monkey supplementary motor area as observed by Berdyyeva and Olson (2010).

**Figure 1:**
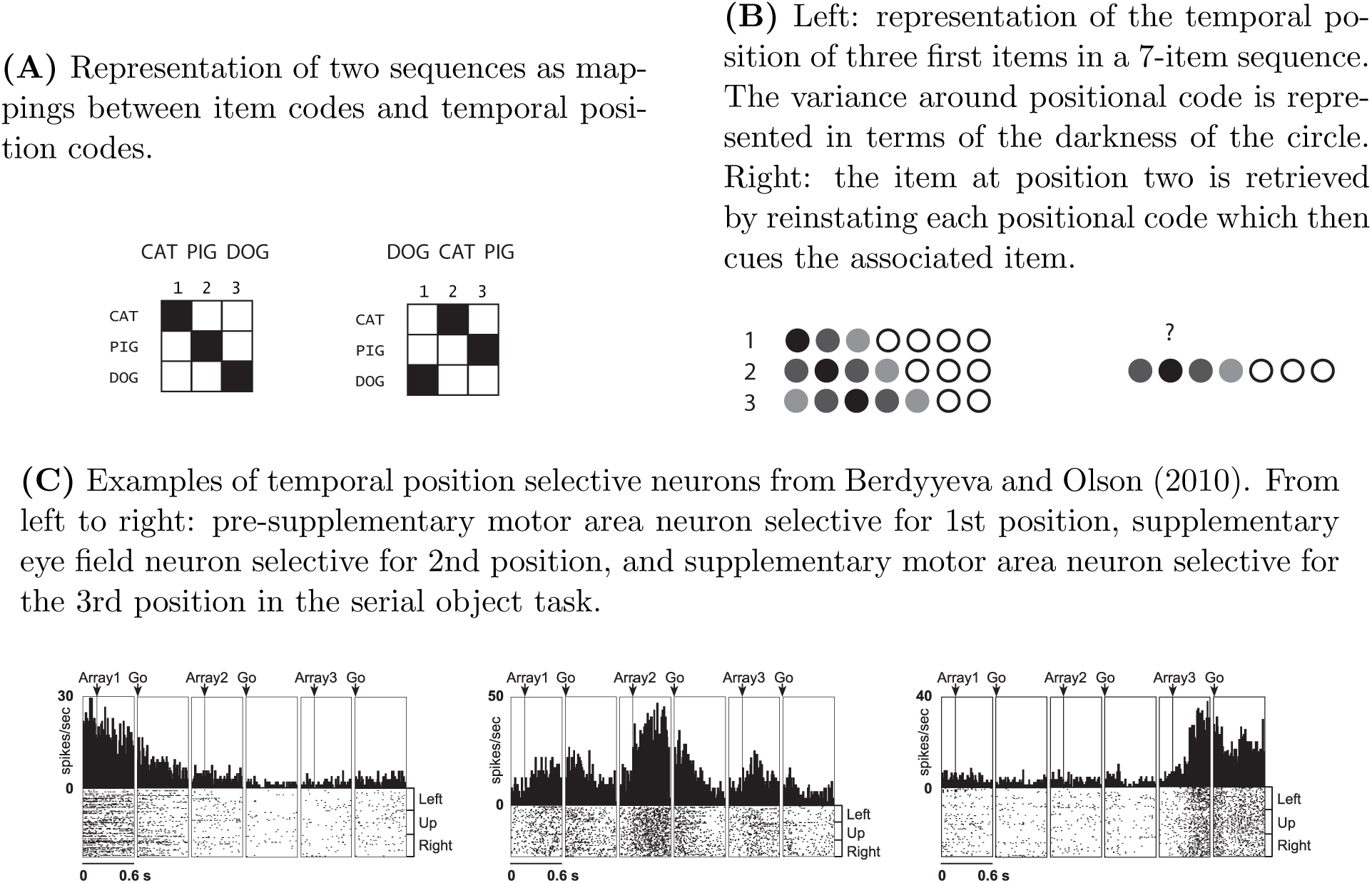
Sequence representation and positional code

Neural implementation of the positional code has been extensively studied with animal neurophysiology. Neurons selective for each position in a sequence have been observed in monkey dorsolateral prefrontal cortex (Averbeck & Lee, 2007; Inoue & Mikami, 2006; Ninokura, Mushiake, & Tanji, 2004; Barone & Joseph, 1989), supplementary and presupplementary motor area (Nakajima, Hosaka, Mushiake, & Tanji, 2009; Berdyyeva & Olson, 2010; Isoda & Tanji, 2004), and medial premotor cortex (Crowe, Zarco, Bartolo, & Merchant, 2014; Merchant, Pérez, Zarco, & Gámez, 2013). Other research on animal and human neurophysiology has suggested that the hippocampus encodes the position of items in a sequence (Heusser, Poeppel, Ezzyat, & Davachi, 2016; Rangel et al., 2014; Hsieh, Gruber, Jenkins, & Ranganath, 2014; DuBrow & Davachi, 2014; Ginther, Walsh, & Ramus, 2011), with some authors proposing the existence of ‘time cells’ tracking the temporal information during sequence processing (MacDonald, Lepage, Eden, & Eichenbaum, 2011; MacDonald, Carrow, Place, & Eichenbaum, 2013).

However, little is known about the neural representation of the positional code in human STM. Previous human neuroimaging studies have instead focussed on learned sequences (Ross, Brown, & Stern, 2009; Albouy et al., 2008; Schendan, Searl, Melrose, & Stern, 2003; Hsieh & Ranganath, 2015; Hsieh et al., 2014) or not used a STM order recall task (Heusser et al., 2016; DuBrow & Davachi, 2016, 2014; Amiez & Petrides, 2007; Hsieh & Ranganath, 2015; Hsieh et al., 2014). Furthermore, several unrelated cognitive processes, such as memory load, sensory adaptation, and reward expectation, also change in a consistent manner as the sequence unfolds. This confound has been thus far ignored by studies using fMRI to investigate the human positional code suggesting that their results must be treated with caution (for a review see Kalm & Norris, 2016).

In the current paper we investigate how the positional code is represented in the brain when a short sequence is recalled after a brief period of time. We used an STM task where participants had to remember and subsequently recognise a short sequence of images. In order to recall items in the correct order participants had to retrieve the positional code instantiated during the presentation phase of the STM task (Figure 1B). Here we investigated whether any brain regions shared this positional code between the presentation and response phases of the task. For this purpose, we trained a classification algorithm to use the fMRI activity patterns of individual items to predict the positions of those items when they appeared in different sequences. Such an analysis abstracts over item identity (such as a specific image) but is sensitive to the representations that consistently code for an item’s position within a sequence. Importantly, we used activity patterns from the presentation phase of the STM task to predict the order of the items during the subsequent recognition phase. This allows us to identify brain regions where the positional code is shared between encoding and response phases and hence relates to memory representations and not other sequential processes such as memory load or sensory adaptation.

Our results reveal that although several brain regions showed sensitivity to order within a single phase only the voxels in the lateral prefrontal cortex (PFC) and the anterior temporal lobe (ATL) represented item position consistently across task phases. This suggests that while many brain areas, including sensory and motor cortices, are sensitive to temporal position, those representations might not be used to guide behaviour and could instead reflect perceptual or load-related aspects of the task. Our findings suggest that voxels in the PFC and ATL are not only sensitive to sequentially presented stimuli (Amiez & Petrides, 2007) or sequentially executed actions (Averbeck & Lee, 2007) but encode temporal position information across task phases in a manner which can be used to guide behaviour.

## 2 Methods

### 2.1 Participants

In total, 13 right-handed volunteers (6 female, 20-33 years old) gave informed, written consent for participation in the study after its nature had been explained to them. Subjects reported no history of psychiatric or neurological disorders and no current use of any psychoactive medications. Two participants were later excluded from the study because of excessive motion artefacts in the collected fMRI data (see Physiological noise removal for the exclusion criteria). The study was approved by the Cambridge Local Research Ethics Committee (LREC) (Cambridge, UK).

### 2.2 Task

We used an immediate serial recognition task where participants had to remember sequences of one, two or three pictures of houses or faces in the order they were presented (Figure 2). On each trial participants were presented with a visual fixation cross to indicate the start of the presentation of the sequence. During the presentation phase, pictures of houses or faces were presented individually (each item for 3.5 seconds) followed by a brief delay (2 seconds). This was followed by an order recognition phase, where a replay of the initial sequence was displayed. At the end of this phase participants had to indicate whether the items in the sequence were presented in the same order as in the original sequence (Figure 2). In order to ensure that participants paid constant attention during recognition we incorporated repeats and omissions in the replayed sequence (8 trials out of 96). Items which did not appear in their original presented positions during those trials were not included in the later fMRI data analysis. On 1/3 of the trials the recognition phase was omitted. The recognition phase was followed by a cue + indicating that there would be a delay of between 6 and 16 seconds before the next trial. The inclusion of the recognition phase in the trial was randomised across the experiment.

**Figure 2:**
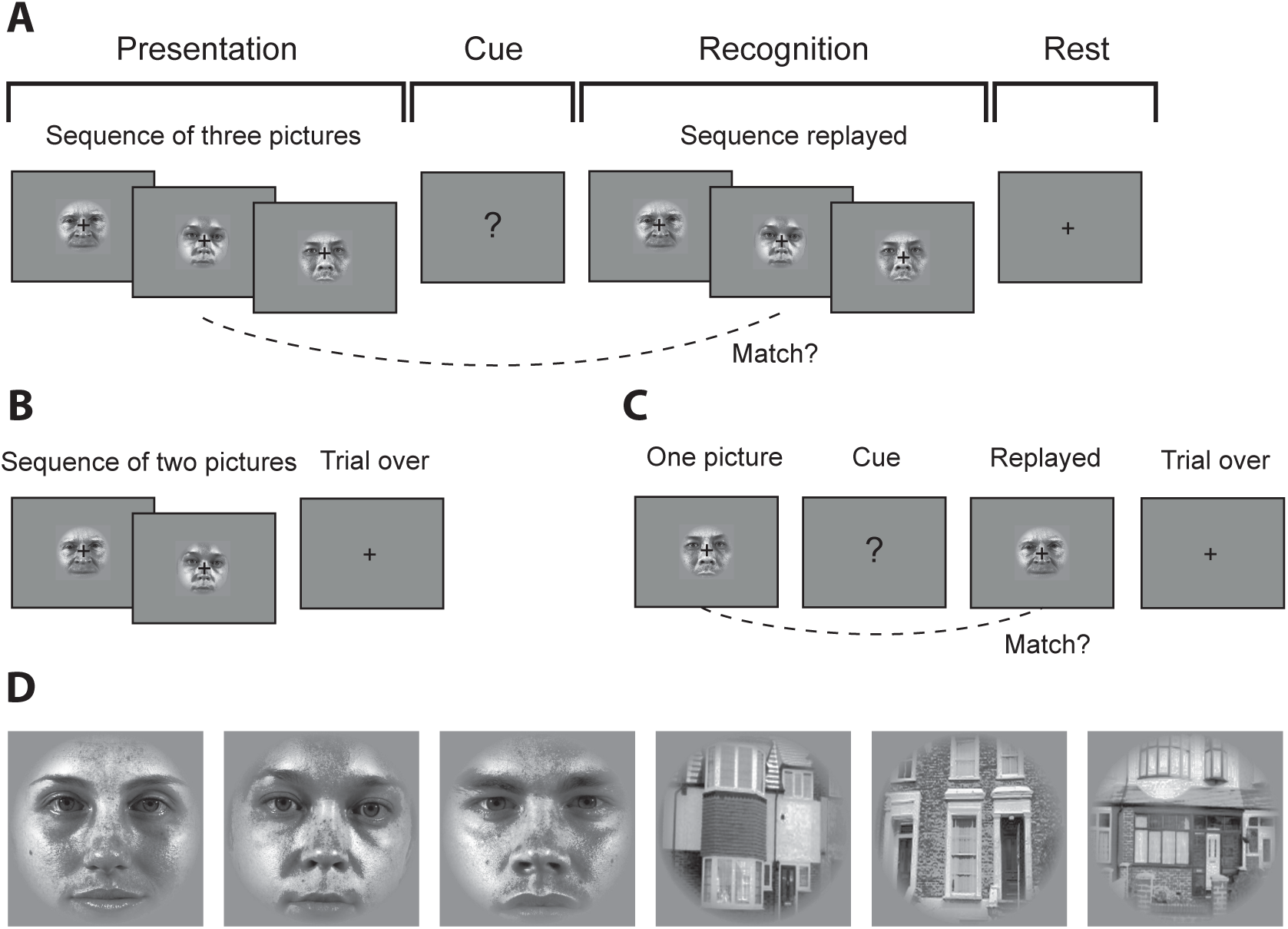
Examples of a single trial: (A) Three-item sequence where all items were presented in the recognition phase, item-order mappings remained the same; (B) two-item sequence without recognition, (C) single item ‘sequence’ with recognition, item-order mapping not the same. (D) Examples of stimuli.

We used short 3-item sequences to ensure that the entire sequence could be accurately retained in STM. If we had used longer sequences then the representation of any given sequence would necessarily vary from trial to trial depending on the nature of the errors, and no consistent pattern of neural activity could be detected. Furthermore, we wanted to estimate separate regressors for individual items in the sequence during both the presentation and recognition phases of the tasks. Presenting stimuli sequentially in an event related fMRI design poses substantial problems for later data analysis: using temporally adjacent stimuli without an intervening rest period creates collinearity in the fMRI data due to the temporal lag of the haemodynamic response function. This in turn makes it difficult to estimate BOLD responses separately for the individual items in the sequence. We took a number of steps to address this issue in the task design. First, the number of items in the sequence was varied randomly across the trials. This ensured that the first item in the sequence was not always followed by a second item, and similarly the second item not by a third. As a result, 44% of the presented sequences were three items long, 39% were two items long and 17% one item long (see Table 1). Second, participants’ memory was probed only on 2/3 of the trials so that we could model the fMRI responses from presentation and response phases separately. Each participant was presented with 96 trials in a single scanning run in addition to an initial practice session outside the scanner. Participants were not informed that there were different types of trials.

**Table 1:**
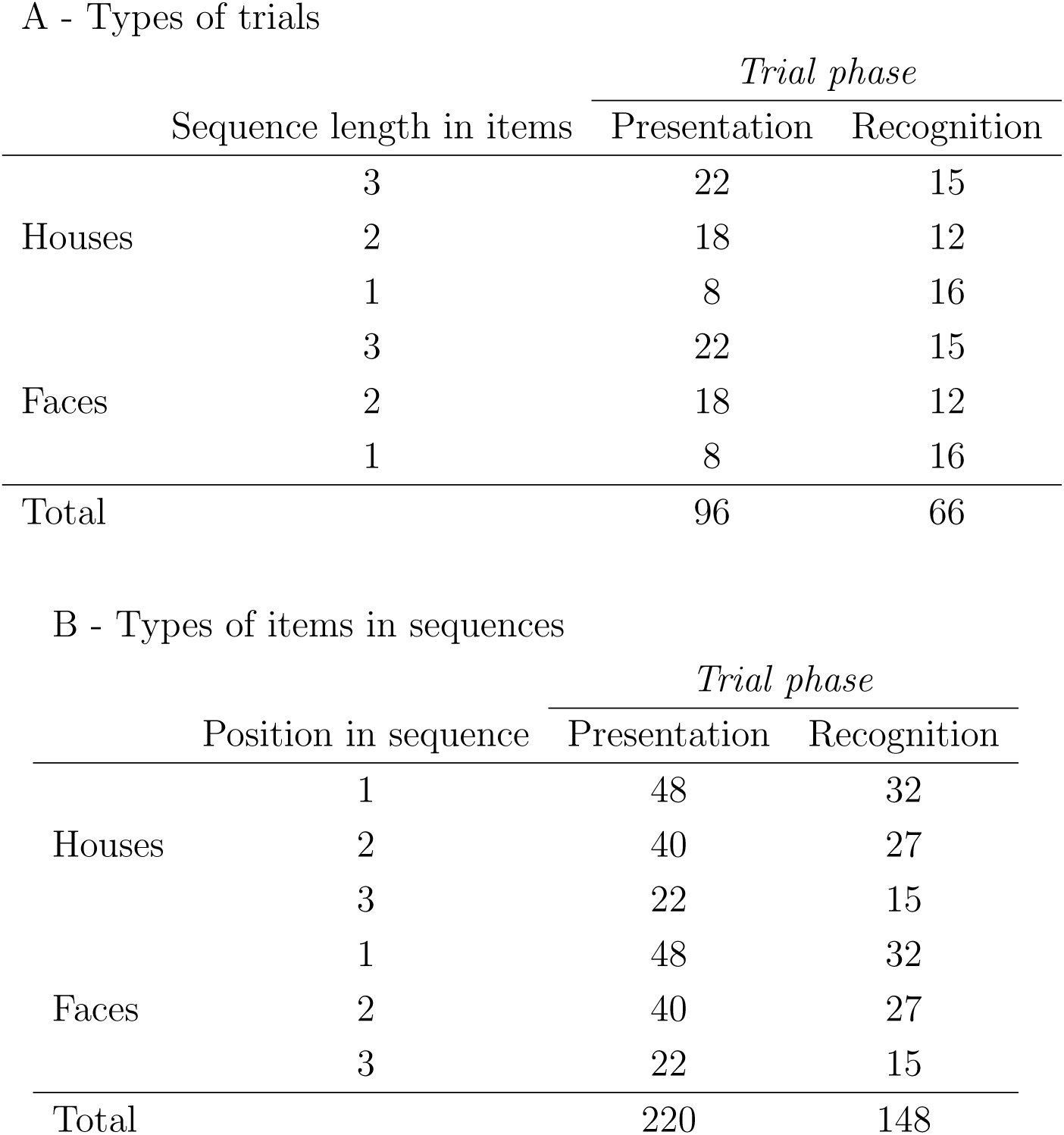
Structure of trials and items presented during the experiment

A - Types of trials
|  |  | *Trial phase* |  |
| --- | --- | --- | --- |
|  | Sequence length in items | Presentation | Recognition |
| Houses | 3 | 22 | 15 |
|  | 2 | 18 | 12 |
|  | 1 | 8 | 16 |
| Faces | 3 | 22 | 15 |
|  | 2 | 18 | 12 |
|  | 1 | 8 | 16 |
| Total |  | 96 | 66 |

B - Types of items in sequences
|  |  | *Trial phase* |  |
| --- | --- | --- | --- |
|  | Position in sequence | Presentation | Recognition |
| Houses | 1 | 48 | 32 |
|  | 2 | 40 | 27 |
|  | 3 | 22 | 15 |
| Faces | 1 | 48 | 32 |
|  | 2 | 40 | 27 |
|  | 3 | 22 | 15 |
| Total |  | 220 | 148 |

### 2.3 Stimuli

The pictures of faces and houses were processed in Matlab to achieve similar luminance histograms, and were cropped to ensure that each image appeared in a similar retinal area. Cropping was achieved with a smooth border, and the resulting image was superimposed on a grey background (Figure 2). The stimuli subtended a 6° visual angle around the fixation point in order to elicit an approximately foveal retinotopic representation. Stimuli were back-projected onto a screen in the scanner which participants viewed via a tilted mirror. The experiment was controlled using Matlab and the Psychophysics Toolbox extension (Kleiner et al., 2007).

### 2.4 fMRI data acquisition and pre-processing

Participants were scanned at the Medical Research Council Cognition and Brain Sciences Unit (Cambridge, UK) on a 3T Siemens TIM Trio MRI scanner using a 32-channel head coil. Functional images were collected using 32 slices covering the whole brain (slice thickness 2mm, 25% slice gap, in plane resolution 2 × 2 mm) with *TR* = 1.75s and *TE* = 44ms. In addition, MPRAGE structural images were acquired at 1mm isotropic resolution. (See http://imaging.mrc-cbu.cam.ac.uk/imaging/ImagingSequences for detailed information.) All volumes were collected in a single, continuous run for each participant. 756 volumes were acquired in a single acquisition run which lasted approximately 22 minutes. The initial six volumes from the run were discarded to allow for T1 equilibration effects. All fMRI data were pre-processed using SPM8 software (Wellcome Trust Centre for Neuroimaging, London) and analysed using custom in-house software. Prior to analysis, all images were corrected for slice timing, with the middle slice in each scan used as a reference. Images were realigned with respect to the first image using tri-linear interpolation, creating a mean realigned image. The mean realigned image was then co-registered with the structural image and the structural image was normalized to the MNI average brain using the combined segmentation/normalization procedure in SPM8. The functional volumes remained unsmoothed and in their native space for participant-specific generalized linear modelling.

### 2.5 Physiological noise removal

In order to remove physiological noise from the fMRI signal we measured respiratory and cardiac data during scanning and used them as nuisance regressors in the general linear model (GLM). A pulse oximeter was used to record participants’ cardiac data, and a pneumatic breathing belt to record the respiratory data. The physiological data were then filtered and down-sampled using the PhLEM toolbox (Verstynen & Deshpande, 2011) to match the scan acquisition time and added to the GLM as separate nuisance regressors. Six motion parameters corresponding to translations and rotations of the image due to movement in the scanner, and additional scan-specific regressors were also added to account for large head movements. Additional parameters were modelled to account for extreme inter-scan movements which exceeded a translation threshold of 0.5mm, rotation threshold of 1.33° and between-images difference threshold of 0.035 calculated by dividing the summed squared difference of consecutive images by the squared global mean. Two participants were excluded from the study because more than 10% of the acquired volumes had extreme inter-scan movements.

### 2.6 General linear model and event regressors

#### 2.6.1 Event regressors

We sought to dissociate fMRI activity patterns representing the identity of the items from the patterns representing their position within the sequence. As noted above, when stimuli are presented in immediate succession without an intervening rest period this creates collinearity in the fMRI data due to the temporal lag of the HRF. We took a number of steps to address this issue in the experiment design (see also Task above). First, we randomised the number of items in the sequence and their order of appearance across the trials. Second, we presented each item for 3.5 seconds to obtain two scans of data per item in the sequence. Third, we omitted the response phase of the task on approximately 1/3 of the trials. Fourth, we jittered the duration of the rest phase between 6-16 seconds. Fifth, no temporal decorrelation or whitening of fMRI data was carried out prior to estimating the general linear model (GLM) to avoid artificial dissimilarities between adjacent events. Finally, every event regressor was estimated with a separate GLM to avoid an overlap in temporally adjacent regressor estimates. This process included combining individual events so that the number of final regressors was balanced across event types (see Table 2).

**Table 2:** Distribution of fMRI data regressors for every participant

|  |  | *Trial phase* |  |
| --- | --- | --- | --- |
|  | Position in sequence | Presentation | Recognition |
| Houses | 1 | 14 | 9 |
|  | 2 | 14 | 9 |
|  | 3 | 14 | 9 |
| Faces | 1 | 14 | 9 |
|  | 2 | 14 | 9 |
|  | 3 | 14 | 9 |
| Total |  | 42 | 27 |

As a result of these measures we obtained a sufficient degree of decorrelation between the event regressors in the GLM for every position in the sequence. Finally, nuisance regressors were added to each GLM modelling head movement and cardiac and respiratory activity (see Physiological noise removal).

#### 2.6.2 GLM estimation

The event regressors were convolved with the canonical haemodynamic response (as defined by SPM8 analysis package) and passed through a high-pass filter (128 seconds) to remove low-frequency noise. Parameter estimates (beta weights) measuring the brain activity evoked during each event type were estimated with the Least-Square2 method (Turner, Mumford, Poldrack, & Ashby, 2012) so that a separate GLM was estimated for each beta. This ensured that the overlap between adjacent event regressors did not affect the estimated beta weights. The resulting beta volumes were grey-matter-masked using the tissue probability maps generated by the segmentation processing stage and were used as inputs for multi-voxel pattern analysis. As a result, we obtained 69 images representing the individual sequence items from both presentation and recognition phases of the experiment (see Table 2) for each participant.

### 2.7 Multi-voxel pattern analysis

A number of methodological issues need to be addressed when performing an fMRI experiment where the aim is to investigate the representation of temporal position. The central problem is that items in different positions necessarily differ along other dimensions too. An item in position three is preceded by more items than one in position two and occurs at a later time than item two. In a memory task, memory load will be greater at position three than position two. Any or all of these factors might lead to an increase or decrease in activation over position, and this would provide the information necessary for a linear classifier to discriminate between items in different positions. These challenges and how they can be overcome are covered in detail by Kalm and Norris (2016). To briefly summarise, we used two methods to ensure that classification was based on the positional code rather than information collinear to the positional information. First, we excluded univariate changes between sequence items by z-scoring the activation of all voxels with respect to their mean activity before the analysis. This ensured that our classification analysis was insensitive to changes which affect a brain region uniformly, such as sensory adaptation or memory load. Second, we employed an analysis where training and testing data came from two different task phases. This ensured that the accuracy of the classification was based on the information shared by two different behavioural stages and hence any effects of the particular task phase (presentation or recognition) would be cancelled out. We refer the reader to Kalm and Norris (2016) for a detailed account of these issues.

#### 2.7.1 Classification analysis

We moved a spherical searchlight with a 6-mm radius throughout the grey-matter masked and unsmoothed volumes to select, at each location, a local contiguous set of 124 voxels (2mm isotropic). As a result, the voxels in the searchlight comprised a vector of activations from beta images resulting in 124 × 69 matrix of *voxels × sequence items*. Voxel vectors where then z-scored to exclude any univariate effects from the analysis.

To identify voxels which encoded position information we ran a 3-way classification of item position in a sequence for both stimulus types (houses and faces). We labelled the voxel vectors according to their position in the sequence (1, 2, or 3; see Stimuli for details) and split the vectors into two data sets: a training set used to train a support vector machine (SVM, with a linear kernel and a regularization hyper-parameter *C* = 40) to assign correct labels to the activation patterns (1, 2 or 3), and a test set (including one sample from each class) used to independently test the classification performance. The SVM classifier was trained to discriminate between the three order positions with the training data, and subsequently tested on the independent test data.

We carried out three classification analyses: position classification during the presentation and recognition phases, and a cross-task phase position classification. In the first two both training and testing data came from the same phase of the task (presentation or recognition). We used leave-3-out cross-validation (one test item per each of 3 classes) to obtain a mean classification accuracy for a searchlight and then calculated an estimate of a true classification accuracy (see Significance testing below). In the cross-phase analysis the training data came from the presentation phase of the trial while the testing data came from the recognition phase of the trial. Here cross-validation was a-priori provided by two task phases. The classification was performed with the LIBSVM (Chang & Lin, 2011) implementation.

For every participant, the classification searchlight analysis resulted in a classification accuracy brain map. We assigned a score of zero to any sphere in which fewer than 33 voxels were inside the individual grey matter volume. These individual images were subsequently normalised to the MNI anatomical template and entered into group-level analyses.

#### 2.7.2 Significance testing

We assessed classification accuracies statistically with non-parametric randomization tests (Stelzer, Chen, & Turner, 2012). We permuted the correspondence between the test labels and data 100 different times to compute 100 mean classification accuracies for the testing labels. To this permuted distribution of accuracies we added the mean accuracy obtained with the correct labelling. We then obtained the distribution of group-level mean accuracies by randomly sampling 1000 mean accuracies (with replacement) from each participant’s permuted distribution. Next, we found the true group-level mean accuracy’s empirical probability based on its place in a rank ordering of this distribution. The peak percentiles of significance (*p* < 0.001) are limited by the number of samples producing the randomized probability distribution at the group level.

## 3 Results

### Behavioural results

All participants performed at a ceiling level or very close to it: the average number of incorrect recognition decisions was 0.6 out of 96 trials (99.4% mean accuracy). This was because short sequences of three or fewer items are easily retained in visual STM (Alvarez & Cavanagh, 2004; Cowan, 2001). The data from incorrectly recalled trials were excluded from the imaging analysis (see Methods, Event regressors).

### Representation of the position of individual items in STM

We ran a whole-brain searchlight classification analysis to identify which brain regions shared the positional code between the presentation and recognition phases of the task. Classification was significantly above chance bilaterally in the rostro-lateral prefrontal cortex (rlPFC) and anterior portion of the superior and middle temporal lobe (Figure 3 – red-yellow accuracy map). There was no significant difference in decoding accuracy between different classes of stimuli (houses and faces, *df* = 12, *p* = 0.63). When classification was carried out within a single task phase only (presentation or recognition) we could decode item position in the sequence within large portions of the lateral occipital cortex and posterior portions of the the temporal lobes (Figure 3 – blue-cyan accuracy map).

**Figure 3:**
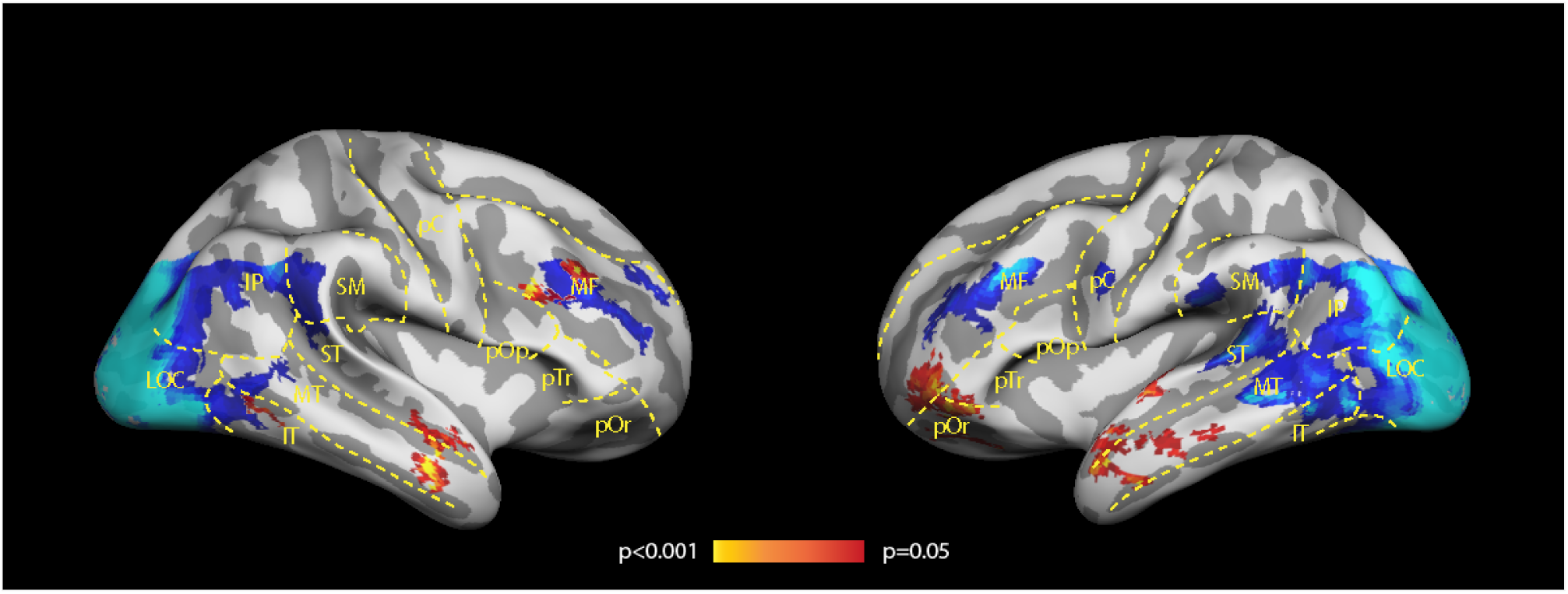
Regions where the position of the items within a sequence was decoded significantly above chance across across participants: red-yellow – significantly above chance between task phases, blue-cyan – significantly above chance within a single task phase only. Abbreviations correspond to the following cortices: MF – middle frontal lobe, pOr – pars orbitalis, pTr – pars triangularis, pOp – pars opercularis, pC – precentral area, ST – superior temporal lobe, MT – middle temporal lobe, IT – inferior temporal lobe, LOC – lateral occipital lobe, SM – supramarginal area.

The analysis of pattern similarity between item positions revealed differences in the way anterio-temporal and prefrontal cortices represent positional code. We observed that in the rlPFC regions the first position in the sequence was significantly misclassified compared to the the second or the third positions. In other words, first position representations in the recognition phase were mostly predicted to be either second or third positions in the sequence. This can be directly observed by comparing the known positions of the items to the predictions made by the classification algorithm (Figure 4A, left column). Contrastingly, in the anterior temporal lobe (ATL) regions all positions were on the average classified correctly (Figure 4B, left column).

**Figure 4:**
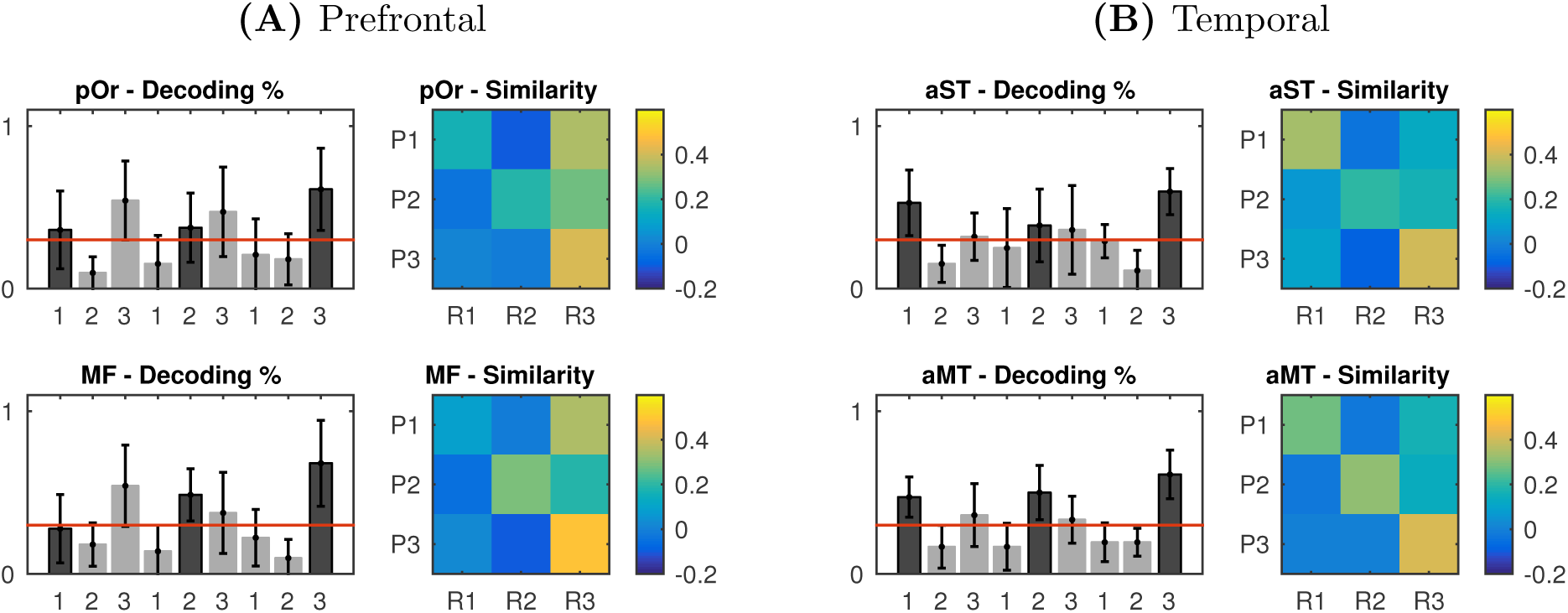
Classification accuracy and pattern similarity between two task phases in prefrontal (A) and temporal regions (B). Bar charts display the average classification accuracy across participants by comparing the known positions (labels) to the predictions made by the classification algorithm. Bars show the proportion of predicted values for each position. Correct classifications are represented with a darker bar. Error bars show the standard error of the mean. The red line depicts the chance level classification accuracy 1/3. Similarity matrices display average pairwise pattern correlations (Pearson’s *ρ*) between two task phases: P – presentation, R – recognition, 1, 2, 3 – position. Cells on the diagonal show the pattern correlation within the same positions between two task phases. Abbreviations: MF – middle frontal lobe, pOr – pars orbitalis, aST – antero-superior temporal lobe, aMT – anterior middle temporal lobe.

These results suggest that in the rlPFC the representation of the first position changed significantly between task phases compared to the representations for the second and third positions. Figure 5A illustrates this with a single subject example: the activation distribution for the first position changed significantly between task phases while the distributions for the second and third positions changed less. Average changes in position representations across subjects (Figure 5B) show that in rlPFC regions the first position in the sequence has consistently changed more than 2nd or 3rd position representations (*df* = 12, *p* < 0.01). Contrastingly this was not true for the ATL regions (Figure 5B, right column).

**Figure 5:**
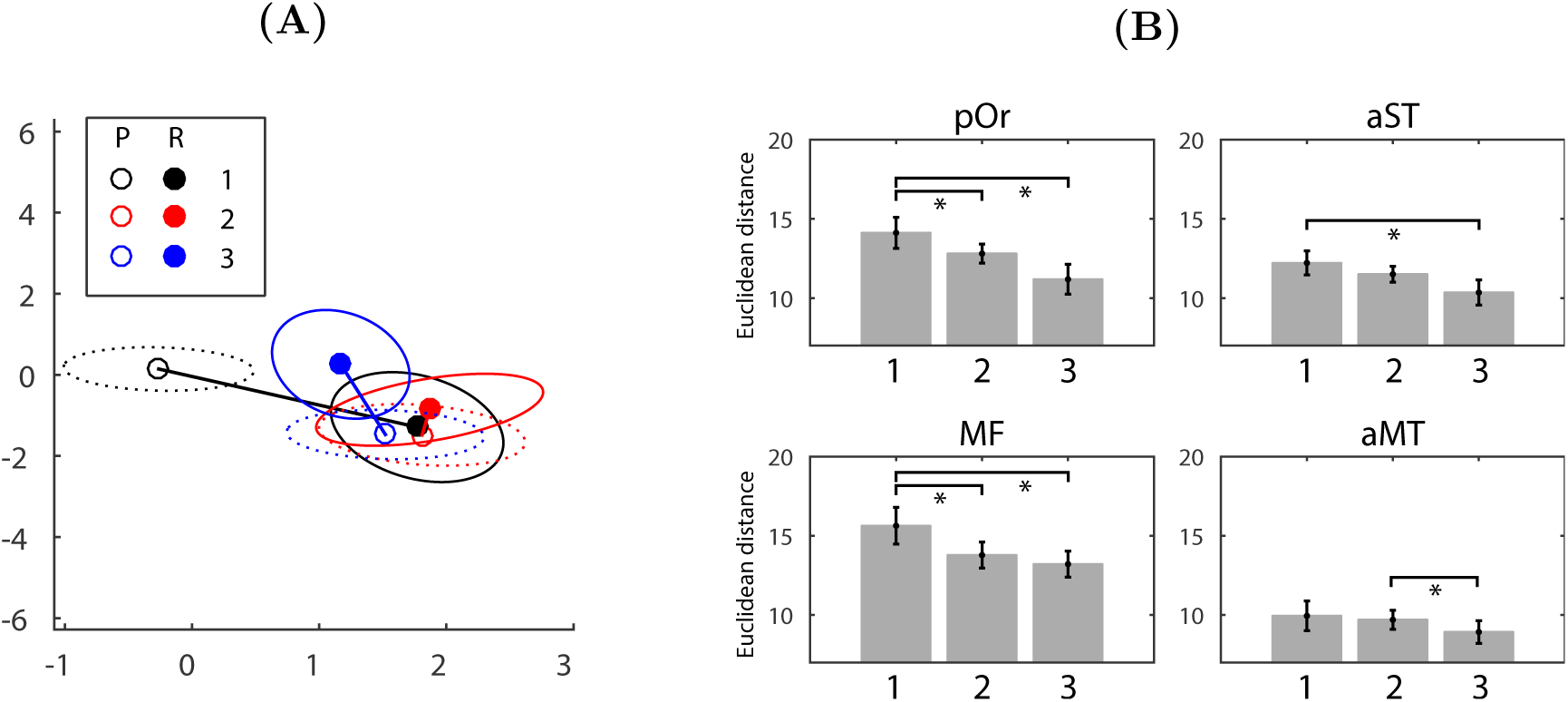
(A) Change in position representations for a single subject between two task phases. For visualisation purposes only values from two most discriminant voxels are plotted. Empty markers represent the distribution of means for the presentation phase, filled markers for the recognition phase. Black – first position, red – second position, blue – third position. Ellipses represent two standard deviations around the mean: dotted ellipses – presentation, solid ellipses – recognition. Straight lines represent the euclidean distance the mean of the distribution has moved in two-voxel space. (B) Change in position representations across task phases averaged over participants. Each bar shows how much the distribution mean for each position representation has moved with respect to the decision boundary of the classifier. Comparisons marked with an asterisk denote *p* < 0.05 differences in means (*df* = 12).

In sum, although we could predict the position of individual items in several brain areas, only regions in the rlPFC and ATL encode position across task phases. However, out of the two regions, only in the ATL the representations for all three sequence positions are consistent over the duration of the STM task: pattern similarity analysis revealed in the rlPFC regions representations for the first positions change so that they could not be reliably decoded during the recognition phase.

## 4 Discussion

In this paper we investigated whether any brain regions shared the positional code between the presentation and response phases of the STM task. We found that while voxels in a number of brain regions represented positional information during either presentation and recognition phases, only voxels in the rostro-lateral prefrontal cortex (rlPFC) and the anterior temporal lobe (ATL) represented position consistently across task phases.

This suggests that a positional ‘read-out’ in the sensory cortices (Figure 3 – blue-cyan accuracy map) is not indicative of STM representations of the positional code but rather reflects perceptual or load-related aspects of the task. Our findings suggest that only position representations in the PFC and ATL encode temporal position information across task phases (Figure 3 – red-yellow). Furthermore, only in the ATL were all three sequence position representations shared between task phases.

### 4.1 The positional code in the lateral ATL

Regions in the lateral ATL have been previously shown to serve linguistic and auditory processing (Visser, Jefferies, & Lambon Ralph, 2010), including semantic features of the stimuli (for a review see Bonner & Price, 2013). This raises a question about the nature of the positional code in the ATL as we used *visual* stimuli in our STM task. Hence, our findings suggest that participants could engage in auditory-verbal recoding of unfamiliar visual stimuli. Numerous studies have observed verbal recoding of visual stimuli in STM tasks (Brandimonte & Gerbino, 1993; Palmer, 2000; Brandimonte, Hitch, & Bishop, 1992). Furthermore, visual sequences have been shown to be encoded differently from auditory sequences, leading to qualitatively different serial position curves (Hurlstone et al., 2014; Crowder, 1986; Conway & Christiansen, 2005). Saffran (2003) showed that repeated presentations of an item in the same position improved learning for auditory stimuli, and for simultaneously presented visual stimuli, but not for sequentially presented visual stimuli. Hence our data, together with previous findings, suggest that unfamiliar visual sequences might be recoded verbally to facilitate the maintenance of positional codes between STM task phases. In other words, as the information about the order of the individual stimuli in the sequence needs to be in active and accessible form in STM over a period of few seconds, verbal recoding and rehearsal might help to retain the positional code between initial instantiation and subsequent recall.

### 4.2 The positional code in the PFC

Our results replicate previous studies which have observed neural positional code in the lateral prefrontal cortices of monkeys (Averbeck & Lee, 2007; Inoue & Mikami, 2006; Ninokura et al., 2004; Barone & Joseph, 1989) and humans. Amiez and Petrides (2007) found that human mid-dorsolateral PFC areas 46 and 9/46 were more activated during the presentation phase of temporal order task compared to the control task of stimulus identification. However, animal studies have almost exclusively used motor tasks to probe temporal order memory by requiring the animal to recreate some aspect of the sequence through motor responses (for a review see Kalm & Norris, 2016). As noted by Averbeck and Lee (2007), this motor component of the task makes it hard to distinguish between sequential action planning and item-independent memory representations of position. Here we show that even without a manual recall task the PFC represents item position between task phases. However, unlike in the ATL regions, not all position representations were shared between the presentation and recognition phases. Specifically, we observed the the first position representation changed significantly after the presentation so that it was consistently misclassified during the recognition phase (Figure 5). This suggests that the positional code in the PFC is either susceptible to processes which evolve along the sequence but do not represent position (such as memory load or sensory adaptation) or represent the start of the sequence in some specific way.

### 4.3 The positional code in the STM task

Our study is the first to examine the position of individual items with fMRI in a STM order task. Previously, Hsieh and Ranganath (2015) carried out two fMRI studies focussing on the representation of learned sequences (Hsieh & Ranganath, 2015; Hsieh et al., 2014). In both studies the authors also presented ‘unlearned’ sequences and analysed the data from those trials in terms of the pattern similarity of the position of individual items. They found that the voxels in the parahippocampal cortex encoded the presented objects in terms of position (Hsieh & Ranganath, 2015; Hsieh et al., 2014). However, neither of these studies investigated STM for order. In both studies participants were not required to retain or recall order information whilst being scanned but instead had to make a semantic judgement on each visually presented object. As a result, it is difficult to suggest that the observed differences in pattern similarities in random sequences were attributable to the representation of a positional code in STM. Contrastingly, here we used a task where participants actively encoded, maintained and recalled an equal number of random sequences within their STM span. This ensured that the position information in each sequence were not yet learned and the representations had to be stored in STM.

Results from physiology and imaging studies in animals and humans indicate that the medio-temporal lobe (MTL) plays a critical role in sequence memory. A large body of evidence suggests that the hippocampus proper encodes the associations between individual items and their positions (Manns, Howard, & Eichenbaum, 2007; Devito & Eichenbaum, 2011; Hsieh et al., 2014; Hsieh & Ranganath, 2015; Heusser et al., 2016; Naya & Suzuki, 2011). When the hippocampus is pharmacologically inactivated, rodents lose the ability to remember the sequential ordering of a series of odours (Devito & Eichenbaum, 2011; Kesner, Hunsaker, & Ziegler, 2010; Fortin, Agster, & Eichenbaum, 2002). In contrast to the studies above, we did not observe shared positional code between the presentation and recognition phases in the hippocampus proper or in anterior parts of the MTL. Although these null results should be interpreted cautiously this could be explained by the relative lack of fMRI signal from the antero-ventral parts of the MTL compared to the more posterior parts of the ventral visual processing stream. However, as noted above, previous studies mostly studied MTL represents of highly learned sequences rather than how that information is maintained in STM.

### 4.4 Conclusions

Our results reveal that only the voxels in the lateral prefrontal cortex and the anterior temporal lobe represented item position consistently across visual STM task phases. This suggests that while many brain areas, including sensory and motor cortices, are sensitive to temporal position, those representations might not be used to guide behaviour and could instead reflect perceptual or load-related aspects of the task. We suggest that shared positional code in the ATL might reflect verbal recoding of visual sequences to facilitate the maintenance of order information over several seconds.

## References

Albouy, G., Sterpenich, V., Balteau, E., Vandewalle, G., Desseilles, M., Dang-Vu, T., … Maquet, P. (2008, apr). Both the hippocampus and striatum are involved in consolidation of motor sequence memory. Neuron, 58(2), 261–72. doi: 10.1016/j.neuron.2008.02.008

Alvarez, G., & Cavanagh, P. (2004, feb). The Capacity of Visual Short-Term Memory is Set Both by Visual Information Load and by Number of Objects. Psychological Science, 15(2), 106–111. doi: 10.1111/j.0963-7214.2004.01502006.x

Amiez, C., & Petrides, M. (2007). Selective involvement of the mid-dorsolateral prefrontal cortex in the coding of the serial order of visual stimuli in working memory. Proceedings of the National Academy of Sciences of the United States of America, 104(34), 13786–91. doi: 10.1073/pnas.0706220104

Averbeck, B., & Lee, D. (2007). Prefrontal neural correlates of memory for sequences. The Journal of Neuroscience, 27(9), 2204–11. doi: 10.1523/JNEUROSCI.4483-06.2007

Baddeley, A. (2003, oct). Working memory: looking back and looking forward. Nature Reviews Neuroscience, 4(10), 829–839. doi: 10.1038/nrn1201

Barone, P., & Joseph, J. (1989). Prefrontal cortex and spatial sequencing in macaque monkey. Experimental brain research, 78(3), 447–64.

Berdyyeva, T., & Olson, C. (2010). Rank signals in four areas of macaque frontal cortex during selection of actions and objects in serial order. Journal of neurophysiology, 104(1), 141–59. doi: 10.1152/jn.00639.2009

Bonner, M., & Price, A. (2013). Where Is the Anterior Temporal Lobe and What Does It Do? Journal of Neuroscience, 33(10), 4213–4215. doi: 10.1523/JNEUROSCI.0041-13.2013

Botvinick, M., & Watanabe, T. (2007). From numerosity to ordinal rank: a gain-field model of serial order representation in cortical working memory. J Neurosci, 27(32), 8636–8642. doi: 10.1523/JNEUROSCI.2110-07.2007

Brandimonte, M., & Gerbino, W. (1993, jan). Mental image reversal and verbal recoding: When ducks become rabbits. Memory & Cognition, 21(1), 23–33. doi: 10.3758/BF03211161

Brandimonte, M., Hitch, G., & Bishop, D. (1992). Influence of short-term memory codes on visual image processing: Evidence from image transformation tasks. Journal of Experimental Psychology: Learning, Memory, and Cognition, 18(1), 157–165. doi: 10.1037/0278-7393.18.1.157

Chang, C., & Lin, C. (2011). LIBSVM. ACM Transactions on Intelligent Systems and Technology, 2(3), 1–27. doi: 10.1145/1961189.1961199

Conway, C., & Christiansen, M. (2005). Modality-Constrained Statistical Learning of Tactile, Visual, and Auditory Sequences. Journal of Experimental Psychology: Learning, Memory, and Cognition, 31(1), 24–39. doi: 10.1016/S0278-7393(05)80004-7

Cowan, N. (2001). The magical number 4 in short-term memory: a reconsideration of mental storage capacity. The Behavioral and brain sciences, 24(1), 87–114; discussion 114–85.

Crowder, R. (1986, apr). Auditory and temporal factors in the modality effect. Journal of experimental psychology. Learning, memory, and cognition, 12(2), 268–78.

Crowe, D., Zarco, W., Bartolo, R., & Merchant, H. (2014). Dynamic Representation of the Temporal and Sequential Structure of Rhythmic Movements in the Primate Medial Premotor Cortex. Journal of Neuroscience, 34(36), 11972–11983. doi: 10.1523/JNEUROSCI.2177-14.2014

Devito, L., & Eichenbaum, H. (2011). Memory for the order of events in specific sequences: contributions of the hippocampus and medial prefrontal cortex. The Journal of Neuroscience, 31(9), 3169–75. doi: 10.1523/JNEUROSCI.4202-10.2011

DuBrow, S., & Davachi, L. (2014). Temporal Memory Is Shaped by Encoding Stability and Intervening Item Reactivation. Journal of Neuroscience, 34(42), 13998–14005. doi: 10.1523/JNEUROSCI.2535-14.2014

DuBrow, S., & Davachi, L. (2016). Temporal binding within and across events. Neurobiology of Learning and Memory. doi: 10.1016/j.nlm.2016.07.011

Fortin, N., Agster, K., & Eichenbaum, H. (2002). Critical role of the hippocampus in memory for sequences of events. Nature Neuroscience, 5(5), 458–62. doi: 10.1038/nn834

Ginther, M., Walsh, D., & Ramus, S. (2011, feb). Hippocampal Neurons Encode Different Episodes in an Overlapping Sequence of Odors Task. Journal of Neuroscience, 31(7), 2706–2711. doi: 10.1523/JNEUROSCI.3413-10.2011

Henson, R., & Burgess, N. (1997). Representations of Serial Order. The Quarterly Journal of Experimental Psychology(789284387).

Henson, R., Norris, D., Page, M., & Baddeley, A. (1996). Unchained Memory: Error Patterns Rule out Chaining Models of Immediate Serial Recall. The Quarterly Journal of Experimental Psychology A, 49(1), 80–115. doi: 10.1080/027249896392810

Heusser, A., Poeppel, D., Ezzyat, Y., & Davachi, L. (2016). Episodic sequence memory is supported by a theta-gamma phase code. Nature neuroscience, 19(August), In Revision. doi: 10.1038/nn.4374

Hsieh, L., Gruber, M., Jenkins, L., & Ranganath, C. (2014). Hippocampal activity patterns carry information about objects in temporal context. Neuron, 81(5), 1165–78. doi: 10.1016/j.neuron.2014.01.015

Hsieh, L., & Ranganath, C. (2015). Cortical and subcortical contributions to sequence retrieval: Schematic coding of temporal context in the neocortical recollection network. NeuroImage, 121, 78–90. doi: 10.1016/j.neuroimage.2015.07.040

Hurlstone, M., Hitch, G., & Baddeley, A. (2014). Memory for serial order across domains: An overview of the literature and directions for future research. Psychological bulletin, 140(2), 339–73. doi: 10.1037/a0034221

Inoue, M., & Mikami, A. (2006). Prefrontal activity during serial probe reproduction task: encoding, mnemonic, and retrieval processes. Journal of neurophysiology, 95(2), 1008–41. doi: 10.1152/jn.00552.2005

Isoda, M., & Tanji, J. (2004). Participation of the primate presupplementary motor area in sequencing multiple saccades. Journal of neurophysiology, 92(1), 653–9. doi: 10.1152/jn.01201.2003

Kalm, K., & Norris, D. (2016, sep). Reading positional codes with fMRI: Problems and solutions. bioRxiv. doi: 10.1101/076554

Kesner, R., Hunsaker, M., & Ziegler, W. (2010). The role of the dorsal CA1 and ventral CA1 in memory for the temporal order of a sequence of odors. Neurobiology of learning and memory, 93(1), 111–6. doi: 10.1016/j.nlm.2009.08.010

Kleiner, M., Brainard, D., Pelli, D., Ingling, A., Murray, R., & Broussard, C. (2007). What’s new in Psychtoolbox-3. Perception 36 ECVP Abstract Supplement.

MacDonald, C., Carrow, S., Place, R., & Eichenbaum, H. (2013). Distinct Hippocampal Time Cell Sequences Represent Odor Memories in Immobilized Rats. Journal of Neuroscience, 33(36), 14607–14616. doi: 10.1523/JNEUROSCI.1537-13.2013

MacDonald, C., Lepage, K., Eden, U., & Eichenbaum, H. (2011, aug). Hippocampal “Time Cells” Bridge the Gap in Memory for Discontiguous Events. Neuron, 71(4), 737–749. doi: 10.1016/j.neuron.2011.07.012

Manns, J., Howard, M., & Eichenbaum, H. (2007). Gradual changes in hippocampal activity support remembering the order of events. Neuron, 56(3), 530–40. doi: 10.1016/j.neuron.2007.08.017

Merchant, H., Pérez, O., Zarco, W., & Gámez, J. (2013, may). Interval tuning in the primate medial premotor cortex as a general timing mechanism. The Journal of neuroscience: the official journal of the Society for Neuroscience, 33(21), 9082–96. doi: 10.1523/JNEUROSCI.5513-12.2013

Nakajima, T., Hosaka, R., Mushiake, H., & Tanji, J. (2009). Covert representation of second-next movement in the pre-supplementary motor area of monkeys. Journal of neurophysiology, 101(4), 1883–9. doi: 10.1152/jn.90636.2008

Naya, Y., & Suzuki, W. (2011). Integrating what and when across the primate medial temporal lobe. Science, 333(6043), 773–6. doi: 10.1126/science.1206773

Ninokura, Y., Mushiake, H., & Tanji, J. (2004). Integration of temporal order and object information in the monkey lateral prefrontal cortex. J Neurophysiol, 91(1), 555–560. doi: 10.1152/jn.00694.2003

Page, M., & Norris, D. (2009, dec). A model linking immediate serial recall, the Hebb repetition effect and the learning of phonological word forms. Philosophical transactions of the Royal Society of London. Series B, Biological sciences, 364(1536), 3737–53. doi: 10.1098/rstb.2009.0173

Palmer, S. (2000, may). Working memory: A developmental study of phonological recoding. Memory, 8(3), 179–193. doi: 10.1080/096582100387597

Rangel, L., Alexander, S., Aimone, J., Wiles, J., Gage, F., Chiba, A., & Quinn, L. (2014). Temporally selective contextual encoding in the dentate gyrus of the hippocampus. Nature communications, 5, 3181. doi: 10.1038/ncomms4181

Ross, R., Brown, T., & Stern, C. (2009, sep). The retrieval of learned sequences engages the hippocampus: Evidence from fMRI. Hippocampus, 19(9), 790–9. doi: 10.1002/hipo.20558

Saffran, J. (2003, aug). Statistical language learning: mechanisms and constraints. Current Directions in Psychological Science, 4(4), 273–114. doi: 10.1111/1467-8721.01243

Schendan, H., Searl, M., Melrose, R., & Stern, C. (2003, mar). An FMRI study of the role of the medial temporal lobe in implicit and explicit sequence learning. Neuron, 37(6), 1013–25. doi: 10.1016/S0896-6273(03)00123-5

Stelzer, J., Chen, Y., & Turner, R. (2012). Statistical inference and multiple testing correction in classification-based multi-voxel pattern analysis (MVPA): Random permutations and cluster size control. NeuroImage. doi: 10.1016/j.neuroimage.2012.09.063

Turner, B., Mumford, J., Poldrack, R., & Ashby, G. (2012). Spatiotemporal activity estimation for multivoxel pattern analysis with rapid event-related designs. NeuroImage, 62(3), 1429–38. doi: 10.1016/j.neuroimage.2012.05.057

Verstynen, T., & Deshpande, V. (2011). Using pulse oximetry to account for high and low frequency physiological artifacts in the BOLD signal. NeuroImage, 55, 1633–1644. doi: 10.1016/j.neuroimage.2010.11.090

Visser, M., Jefferies, E., & Lambon Ralph, M. (2010). Semantic processing in the anterior temporal lobes: a meta-analysis of the functional neuroimaging literature. Journal of cognitive neuroscience, 22(6), 1083–94. doi: 10.1162/jocn.2009.21309

